# Reconstructing Austronesian population history in Island Southeast Asia

**DOI:** 10.1101/005603

**Authors:** Mark Lipson, Po-Ru Loh, Nick Patterson, Priya Moorjani, Ying-Chin Ko, Mark Stoneking, Bonnie Berger, David Reich

## Abstract

Austronesian languages are spread across half the globe, from Easter Island to Madagascar. Evidence from linguistics and archaeology indicates that the “Austronesian expansion,” which began 4–5 thousand years ago, likely had roots in Taiwan, but the ancestry of present-day Austronesian-speaking populations remains controversial. Here, focusing primarily on Island Southeast Asia, we analyze genome-wide data from 56 populations using new methods for tracing ancestral gene flow. We show that all sampled Austronesian groups harbor ancestry that is more closely related to aboriginal Taiwanese than to any present-day mainland population. Surprisingly, western Island Southeast Asian populations have also inherited ancestry from a source nested within the variation of present-day populations speaking Austro-Asiatic languages, which have historically been nearly exclusive to the mainland. Thus, either there was once a substantial Austro-Asiatic presence in Island Southeast Asia, or Austronesian speakers migrated to and through the mainland, admixing there before continuing to western Indonesia.

---

The history of the Austronesian (AN) expansion and of populations speaking AN languages has long been of interest. Patterns of lexical diversity within the AN language family point to Taiwan as the AN homeland (1, 2), as do elements of the archaeological record, for example red-slipped pottery and Taiwanese-mined nephrite (3–5). However, some authors have argued that the AN expansion was driven primarily by cultural diffusion rather than large-scale migration (6–8), and other associated artifacts, such as cord-marked and circle-stamped pottery, likely derive instead from the mainland (9, 10). It is also unknown how the history of populations in western Island Southeast Asia (ISEA), which speak Western Malayo-Polynesian AN languages, differs from that of Central and Eastern Malayo-Polynesian speakers in eastern Indonesia and Oceania.

Genetic data can be used to trace human migrations and interactions in a way that is complementary to the information provided by linguistics and archaeology. Some single-locus genetic studies have found affinities between Oceanian populations and aboriginal Taiwanese (11–15), but others have proposed that present-day AN speakers do not have significant genetic inheritance from Taiwan (16–18). Within Indonesia, several surveys have noted an east–west genetic divide, with western populations tracing a substantial proportion of their ancestry to a source that diverged from Taiwanese lineages 10–30 thousand years ago (kya), which has been hypothesized to reflect a pre-Neolithic migration from Mainland Southeast Asia (MSEA) (19–22). Genome-wide studies of AN-speaking populations, which in principle can provide greater resolution, have been interpreted as supporting both Taiwan-centered (23, 24) and multiple-wave (21) models. However, such work has relied primarily on clustering methods and fitting bifurcating trees that do not model historical admixture events, even though it is well known that many AN-speaking populations are admixed (21, 24–28). Thus, these studies have not established firmly whether AN speakers have ancestry that is descended from Taiwan, MSEA, or both.

Here, we explore these questions by reconstructing the genome-wide ancestry of a diverse sample of AN-speaking populations, predominantly within ISEA. We apply novel methods for determining the phylogenetic placement of sources of gene flow in admixed populations and identify four major ancestry components, including one linked to Taiwan and a second Asian component from MSEA.

## Results

### Analysis of admixed populations

To investigate the ancestry of AN-speaking populations at high resolution, we analyzed a genome-wide data set of 31 AN-speaking and 25 other groups from the HUGO Pan-Asian SNP Consortium (25) and the CEPH-Human Genome Diversity Panel (HGDP) (29). We used genotypes from 18,412 single nucleotide polymorphisms (SNPs) that overlapped across all samples (see Methods, Supplementary Table 1, and Supplementary Fig. 1). To confirm that our results are robust to the way SNPs were chosen, we repeated our primary analyses with data obtained by merging the Pan-Asia genotypes with HGDP samples typed on the Affymetrix Human Origins array (30) (see Methods and Supplementary Tables 6 and 7). For some tests requiring denser markers, we also used a smaller set of 10 AN-speaking groups first published in ref. (27) and typed at over 500,000 SNPs.

We developed new methods to analyze the data, which we release here as the *MixMapper* 2.0 software. *MixMapper* is a tool for building phylogenetic models of population relationships that incorporate the possibility of admixture. Both the original version (31) and *MixMapper* 2.0 use allele frequency correlations to construct an unadmixed scaffold tree and then add admixed populations. The entire best-fitting model for each admixed population, including mixture proportions and the placement of the sources of ancestry on the scaffold, is inferred from the data, and uncertainty in parameter estimates is measured through bootstrap resampling (see Methods). *MixMapper* 2.0 substantially improves the three-way mixture fitting procedure of the original program, as it implements a rigorous test to determine whether populations are best modeled via two- or three-way admixtures. It also allows for full optimization of the inferred mixture proportions (see Methods). A strength of *MixMapper* and related methods is that the underlying allele frequency correlation statistics, and hence the inferences about population relationships, are largely robust to the way that SNPs are chosen for analysis (30–32).

We selected a scaffold tree consisting of 18 populations that are approximately unadmixed relative to each other (Fig. 1; Supplementary Tables 2 and 3): Ami and Atayal (aboriginal Taiwanese); Miao, She, Jiamao, Lahu, Wa, Yi, and Naxi (Chinese); Hmong, Plang, H’tin, and Palaung (from Thailand); Karitiana and Suruí (South Americans); Papuan (from New Guinea); and Mandenka and Yoruba (Africans). This set was designed to include a diverse geographical and linguistic sampling of Southeast Asia (in particular Thailand and southern China) along with outgroups from other continents, which are necessary for accurate mixture fitting (31) (see Methods). We have previously shown that *MixMapper* results are robust to the choice of scaffold populations (31), and indeed our findings here were essentially unchanged when we repeated our analyses with an alternative, 15-population scaffold (Supplementary Fig. 2; Supplementary Tables 8 and 9) and with 17 perturbed versions of the original scaffold (Supplementary Tables 10 and 11). Using this scaffold tree, we obtained confident results for 25 AN-speaking populations (for geographical locations, see Fig. 2): eight from the Philippines, nine from eastern Indonesia and Oceania, and eight from western ISEA. Several populations in our data set—Batak Karo, Ilocano, Malay, Malay Minangkabau, Mentawai, and Temuan—were not as readily fit with *MixMapper*, which we hypothesize was due to the presence of additional ancestry components that we could not capture well in our modeling framework. Thus, we omit these populations from further analyses, although we note that their *MixMapper* results, while not as reliable, were still similar to those for the 25 groups discussed here.

**Figure 1.**
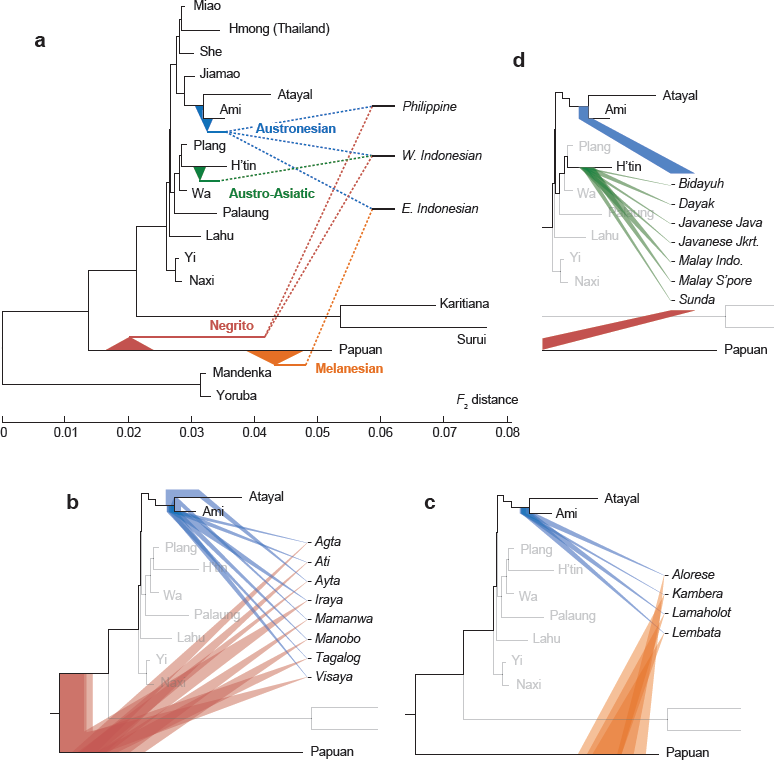
Inferred sources of ancestry for selected admixed Austronesian-speaking populations. Shaded ranges represent 95% bootstrap confidence intervals for branching positions; see Supplementary Tables 4 and 5 for complete mixing branch distributions. The topology of the scaffold tree is shown using the full data set (slight variations are possible across bootstrap replicates). (A) Overview of the three best-fitting admixture models. (B)–(D) Detailed results for highest-confidence models of populations from (B) the Philippines, (C) eastern Indonesia, and (D) western ISEA. In (D), the Austronesian and Negrito branch positions are fixed in *MixMapper* to equal those for Manobo. Batak Toba are omitted for display purposes, as 8% of replicates place their third ancestry component on a non-adjacent branch in the scaffold (Supplementary Table 5). Three other populations (Manggarai Ngada, Manggarai Rampasasa, and Toraja) fall into an additional category of three-way admixed eastern Indonesians, while Oceanians (Fiji and Polynesia) are inferred to have similar ancestry to the populations in (C), but their confidence intervals are not directly comparable because they have fewer SNPs available (see Fig. 2 and Supplementary Tables 4 and 5).

**Figure 2.**
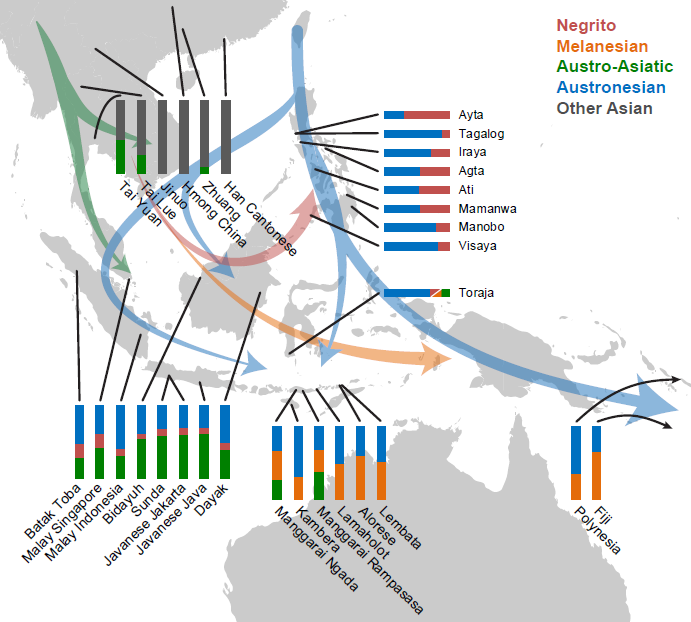
Locations and best-fit mixture proportions (see Methods) for Austronesian-speaking and other populations, with possible directions of human migrations supported by our analyses. For Toraja, we could not distinguish between Negrito and Melanesian ancestry and show this component as red/orange.

All admixed AN-speaking populations fit best as combinations of two or three ancestry components out of a set of four: one closely related to Papuans (“Melanesian”), one splitting deeply from the Papuan branch (“Negrito”), one most closely related to aboriginal Taiwanese, and one most closely related to H’tin (Fig. 1). While the relative proportions varied substantially from group to group, the (independently inferred) positions of the ancestral mixing populations were highly consistent, leading us to assign them to these four discrete sources (Fig. 1). A total of 14 populations were best modeled as two-way admixed (Supplementary Table 4): all eight from the Philippines (with Taiwan-related and Negrito ancestry), four from eastern Indonesia (with Taiwan-related and Melanesian ancestry), and both from Oceania (Fiji and Polynesia, merged from ref. (27); also Taiwan-related and Melanesian). The remaining 11 populations, including all eight from western ISEA, fit best as three-way admixed (Supplementary Table 5), with both Taiwan-related and H’tin-related ancestry (Supplementary Table 12). Among the 25 groups, the Taiwan-related component was inferred to account for approximately 30–90% of ancestry, while for the 11 three-way admixed groups, the H’tin-related component was inferred to account for approximately 10–60%. By contrast, we found no Taiwan-related ancestry in admixed MSEA populations speaking non-AN languages (Fig. 2; Supplementary Table 13). We note that our estimates of mixture proportions are robust to alternative histories involving multiple waves of admixture or continuous migration, since *MixMapper* is based on allele-sharing statistics that measure the probability of descent from each possible source of ancestry. Thus, continuous gene flow scenarios that preserve the same topology relating the admixed population to the scaffold tree will produce the same estimates of mixture proportions (30, 31).

To obtain an independent estimate of how many sources of admixture are necessary to explain the observed relationships among populations from ISEA, we applied a formal test (33,34) that analyzes *f*_4_ statistics among a set of admixed and outgroup populations to determine a lower bound on the total number of ancestry sources (Supplementary Table 14). For the Philippines, we found that a maximal subset of six groups (Agta, Ati, Ayta, Ilocano, Iraya, and Manobo) could be consistently modeled as derived from a single pair of mixing populations (Supplementary Fig. 1A). Likewise, the four eastern Indonesian groups (Alorese, Kambera, Lamaholot, and Lembata) that were inferred to be two-way admixed by *MixMapper* could be modeled with two total ancestry sources according to the *f*_4_-based test (Supplementary Fig. 1B). However, adding the two Manggarai populations required a third source of ancestry, consistent with the H’tin-related ancestry inferred by *MixMapper*. In western ISEA, a large subset of six groups (Bidayuh, Dayak, Javanese Jakarta, Javanese Java, Mentawai, and Sunda) was consistent with being derived from three ancestral mixing populations (Supplementary Fig. 1C), and moderately diverged subsets with as few as three populations (Bidayuh, Dayak, and either Javanese or Sunda) still required three sources of ancestry. Larger subsets were always of greater complexity, indicating some additional, more localized gene flow, such as a likely influx of Indian ancestry in some populations (20, 25). However, the presence of the subsets that can be fit as mixtures of two or three sources increases our confidence that the *MixMapper* models are close to the true history.

Finally, we used our recently developed ALDER software (35) to estimate dates of admixture using linkage disequilibrium. For populations from the Philippines, eastern Indonesia, and Oceania from ref. (27), we obtained dates of 30–65 generations ago assuming a single-pulse model of admixture (0.9–1.8 kya assuming 29 years per generation (36); Supplementary Fig. 3). These dates are considerably more recent than the initial AN expansion as documented through archaeology (2–5), and thus they must reflect additional waves of interaction involving populations with different proportions of Asian ancestry after the initial AN settlement of the islands. We also applied *ALDER* to a merged set of populations from western ISEA and estimated that their admixture occurred 76 ± 21 generations ago (2.2 ± 0.6 kya; Supplementary Fig. 4). Again, this date implies the most recent possible time for the onset of population mixing and should not be interpreted as an estimate of the date of the earliest episodes of admixture (35).

### Details of inferred ancestry components

Our results indicate that there is a component of ancestry that is universal among and unique to AN speakers and that always accounts for at least a quarter of their genetic material. This component, moreover, is more closely related to aboriginal Taiwanese than to any population from the mainland. In theory, this ancestry could have been derived from a mainland source that was related to the ancestors of aboriginal Taiwanese but was either displaced by subsequent migrations (such as the expansion of Han Chinese) or whose descendants are not included in our data set. Given our dense sampling of East and Southeast Asian populations, this scenario seems unlikely, but we are unable to formally rule it out.

We also considered the possibility that the direction of flow for this “Austronesian” ancestry component could have been reversed, with an origin in Indonesia or the Philippines and a northward spread to Taiwan. Because of migrations, it is impossible to determine with certainty where ancestral populations lived based on present-day samples, but the fact that the aboriginal Taiwanese populations in our data set, Ami and Atayal, are unadmixed (to within the limits of our resolution), whereas the AN component appears in admixed form in all other AN-speaking populations from ISEA, can be most parsimoniously explained by a Taiwan-to-ISEA direction of gene flow. We verified that Ami and Atayal have no detectable signature of admixture both by the three-population test (30, 37) (Supplementary Table 3) and by testing them as putatively admixed in *MixMapper* with a scaffold tree made up of the other 16 original scaffold populations. In the latter analysis, we found that both Ami and Atayal returned best-fitting positions that indicated that they are properly modeled as unadmixed, adjacent to Jiamao (Supplementary Table 15). On the other hand, all other AN-speaking populations, including those with no signal of admixture from the three-population test, continued to fit robustly as admixed on this reduced scaffold, with the AN component now closest to Jiamao, as expected (Supplementary Table 15). Thus, the absence of admixture in Ami and Atayal allows us to conclude that they have a qualitatively different history from other AN-speaking populations in ISEA and that our inferred directionality of gene flow, with Taiwan as the source, is more parsimonious and a better fit to the data.

The second and third ancestry components we infer for AN-speaking populations are Melanesian and Negrito. All admixed groups we tested contain at least one of these components, which we believe reflect admixture with indigenous populations in ISEA. The Melanesian component is closely related to Papuans and is found in the highest proportions among our study populations in easternmost Indonesia and in Fiji (Fig. 2). The Negrito component, meanwhile, forms a deep clade with Papuans and is found in populations from the Philippines and western ISEA (Fig. 2). We treat this ancestry as deriving from a single ancient source because it clusters phylogenetically across admixed populations, with the branching positions from the scaffold tree inferred to be very similar (Fig. 1B). We use the name “Negrito” to describe this ancestry based on the fact that it occurs in the greatest proportion in Philippine Negrito populations. The Negrito ancestry in western ISEA could be a result of admixture with aboriginal peoples living on these islands or alternatively of prior admixture in the Philippines or on the mainland. We note that with *MixMapper*, we are unable to determine the precise branching position of this component in three-way admixed populations (see Methods), which would in principle shed light on this question. We are also unable to rule out a small proportion of Negrito ancestry in eastern Indonesia and Oceania—which might be plausible if AN speakers migrated from Taiwan through the Philippines first and admixed at that time with indigenous peoples—or a small proportion of Melanesian ancestry in the Philippines, but the large genetic drift separating the branching positions of the two components (Melanesian and Negrito) provides strong evidence that they reflect at least two ancestral sources (Fig. 1).

An unanticipated finding from our study is that populations in western ISEA, as well as a few in eastern Indonesia, also contain an unambiguous signal of an additional source of Asian ancestry, which is assigned with high confidence to an ancestral population splitting roughly two-fifths of the way down the H’tin branch in our scaffold tree (Fig. 1D). The H’tin speak a language belonging to the Austro-Asiatic (AA) family, which is hypothesized to have been the major language group in MSEA following the expansion of rice farming (5). Later dispersals have resulted in substantial replacements of AA languages outside of Cambodia and Vietnam, but AA-speaking tribal groups are still present in areas where Tai, Hmong, and Indo-European languages now predominate, extending as far west as India (5). By contrast, no pockets of AA languages are found at all in present-day ISEA (with the exception of the Nicobar Islands in the Indian Ocean), which, in conjunction with the absence of clear archaeological evidence of previous settlement by agriculturalists who were not part of the AN cultural complex (10), makes it unlikely that AA-speaking populations previously lived in the areas where we detect AA-related ancestry.

To test the alternative explanation that the genetic evidence of AA-related ancestry in AN speakers might be an artifact of a back-migration from ISEA that contributed ancestry to the H’tin, we removed H’tin from our scaffold tree and repeated our analysis for three-way admixed populations. We found that the formerly H’tin-related ancestry component is now confidently inferred to form a clade with Plang (primarily) or Wa, both of which speak AA languages (Supplementary Table 16). Similarly, when we also removed Plang, it formed a clade with Wa (Supplementary Table 16). We also applied *MixMapper* to two admixed Negrito populations (Jehai and Kensiu) from peninsular Malaysia and found that their Asian ancestry component branches closest to H’tin, in almost exactly the same location as the H’tin-related component from ISEA. Since the Jehai and Kensiu speak AA languages, it is likely that the population contributing their Asian ancestry did as well, and AA-related populations may once have been more widespread in this region. We conclude that our signal indeed reflects gene flow from the mainland into ISEA from an ancestral population that is nested within the radiation of AA-speaking populations, and hence it is likely that this source population itself spoke an AA language.

## Discussion

While a major AA contribution to western speakers of AN languages has not been proposed in the genetic literature, results from previous genetic studies are in fact consistent with these findings. A clustering analysis of the Pan-Asia SNP data (25) showed a component of ancestry in populations from (primarily western) ISEA that also appeared in AA speakers on the mainland, and a separate study of the same data also related western ISEA ancestry to mainland sources (21). However, neither analysis concluded that these signals reflected an AA affinity. Our results are also compatible with published analyses of mtDNA and Y chromosomes, which have provided evidence of a component of ancestry in western but not eastern ISEA that is of Asian origin (20–22). The O-M95 Y-chromosome haplogroup, in particular, is prevalent in western Indonesia (20) and was previously linked to AA-speaking populations (38).

A potential explanation for our detection of AA ancestry in ISEA is that a western stream of AN migrants encountered and mixed with AA speakers in Vietnam or peninsular Malaysia, and it was this mixed population that then settled western Indonesia (Fig. 2). This scenario is consistent with the AN mastery of seafaring technology and would be analogous to the spread of populations of mixed AN and Melanesian ancestry from Near Oceania into Polynesia (13, 15). Since we are unable to determine the date of initial AN–AA admixture, and genetic data from present-day populations do not provide direct information about where historical mixtures occurred, other scenarios are also conceivable; in particular, we cannot formally rule out a wider AA presence in ISEA before the AN expansion or a later diffusion of AA speakers into western ISEA. However, the absence of AA languages in Indonesia, together with our observation of *both* AA and AN ancestry in all surveyed western ISEA populations, suggests that the admixture took place before either group had widely settled the region. We note that in its simplest form, the model of a single early admixture event would imply that populations today should have equal proportions of AN and AA ancestry, which is not the case for our sampled groups. However, these differences could have arisen through a number of straightforward demographic processes, including settlement of different islands by populations with different ancestry proportions, independent fluctuations within populations having heterogeneous ancestry soon after admixture, or continuous or multiple-wave gene flow over a number of generations. Overall, the uniformity of ancestry observed today, with the same components present in all of our sampled groups from western ISEA, points toward a shared mixture event rather than separate events for each population.

These results show that the AN expansion was not solely a process of cultural diffusion but involved substantial human migrations. The primary movement, reflected today in the universally-present AN ancestry component, involved AN speakers from an ancestral population that is most closely related to present-day aboriginal Taiwanese. In western ISEA, we also find an Asian ancestry component that is unambiguously nested within the variation of present-day AA speakers, which makes it likely that the ancestral population itself spoke an AA language. Other suggestions of AN–AA interaction come from linguistics and archaeology (9), as Bornean AN languages contain probable AA loan words (7), and there is evidence that rice (3, 6, 7, 10) and taro (7) cultivation, as well as domesticated pigs (39), were introduced from the mainland. Interestingly, all languages spoken today in both eastern and western ISEA are part of the AN family, which raises the question of why AN languages were always retained by admixed populations. An important direction for future work is to increase the density of sampling of populations from Southeast Asia, with larger sample sizes and more SNPs, if possible in conjunction with ancient DNA (40), to allow more detailed investigation of the dates and locations of the admixture events we have identified.

## Methods

### Data set assembly

For our primary analyses, we merged data from the HUGO Pan-Asian SNP Consortium (25) and the CEPH-Human Genome Diversity Panel (HGDP) (29), yielding a set of 1,094 individuals from 56 populations typed at 18,412 overlapping SNPs. We excluded likely duplicate samples, twins, and first-degree relatives from the Pan-Asia data (a total of 79 individuals) as identified in ref. (41). We also removed 27 individuals identified as outliers by projecting each population onto principal components using EIGENSOFT (42) and deleting samples at least 5 standard deviations away from the population mean on any of the first three PCs.

We also used 10 populations from ref. (27), from a version of the published data set merged with HapMap3 populations but not with Neanderthal and Denisova, for a total of 564,361 SNPs. We restricted to these populations when running *ALDER* and used all of the SNPs. We also merged these samples with our primary data set, leaving 7,668 SNPs, in order to estimate *MixMapper* parameters for Polynesia and Fiji.

In order to test robustness to SNP ascertainment, we repeated our *MixMapper* analyses with a data set formed by merging the Pan-Asia data with HGDP samples typed on the Affymetrix Human Origins array (30), replicating our primary data set on a different collection of 9,032 SNPs. Importantly, the Human Origins SNPs are chosen according to a very different strategy, having been selected based on their presence as heterozygous sites in sequenced genomes from diverse individuals.

Full details for all analyzed populations can be found in Supplementary Table 1.

### Admixture inference with *MixMapper*

The *MixMapper* software estimates admixture parameters using allele frequency moment statistics under a tree-based instantaneous admixture model (31). The program works in two phases. First, it constructs an (approximately) unadmixed scaffold tree via neighbor-joining on a subset of populations chosen by the user to have a specified level of geographic coverage with minimal evidence of admixture based on *f*-statistics (30, 37). The selection of populations for the scaffold is guided by running the 3-population test (30,37), which removes clearly admixed populations; by testing the additivity of possible subtrees from among the remaining populations (similar to the 4-population test (30,37)); and finally by comparing the fits of closely related candidate populations when modeled as admixed. After the scaffold is chosen, the software finds the best-fitting parameters for admixed populations by solving a system of moment equations in terms of the pairwise distance measure *f*_2_, which is the expected squared allele frequency difference between two populations. Specifically, the distance *f*_2_(*C, X*) between an admixed population *C* and each population *X* on the scaffold tree can be expressed as an algebraic combination of known branch lengths along with four unknown mixture parameters: the locations of the split points of the two ancestral mixing populations from the scaffold tree, the combined terminal branch length, and the mixture fraction *α*. In this way, the entire tree topology can be determined automatically, even for large numbers of populations. Finally, *MixMapper* uses a non-parametric bootstrap (43) to determine confidence intervals for the parameter estimates, dividing the SNPs into 50 blocks and resampling the blocks at random with replacement for each of 500 replicates. We note that the bootstrap is applied over the entire fitting procedure, including the application of neighbor-joining to build the scaffold, so that uncertainty in the scaffold topology is accounted for in the final confidence intervals.

For our analyses here, we developed new inference algorithms, released in the *MixMapper* 2.0 software, which extend the original *MixMapper* three-way mixture-fitting procedure, whereby one ancestral mixing population is taken to be related to a population already fit by the program as admixed. First, *MixMapper* 2.0 implements a method to determine the best fit among alternative admixture models—namely, fitting a test population *C* either as two-way admixed or as three-way admixed with one ancestor related to a fixed admixed population *A* (for our applications, either Manobo or Alorese)—by comparing the norm of the vector of residual errors for all pairwise distances *f*_2_(*C, X*), where X ranges over the scaffold populations. Importantly, the two models have the same number of degrees of freedom, with four parameters being optimized in each case. Also, the comparison is restricted those populations *X* on the initial scaffold, i.e., we do not include *f*_2_(*C, A*) in the vector of residuals for the three-way model. Thus, our procedure is conceptually equivalent to augmenting the scaffold by adding *A* (via the standard *MixMapper* admixture model) and then finding the best-fitting placement for *C*. Second, for populations that are better fit as three-way admixed, *MixMapper* 2.0 implements improved estimation of their proportions of ancestry from all three components by re-optimizing this same set of equations but now allowing all of the mixture fractions to vary (as well as the terminal branch lengths for the admixtures, since these depend on the mixture fractions (31)). To prevent overfitting, we fix the branching positions of each ancestry component as determined from the initial fit (independently for each bootstrap replicate).

## Acknowledgments

We thank Peter Bellwood, Nicole Boivin, Richard Meadow, and Michael Witzel for comments on the manuscript. M.L. and P.L. acknowledge NSF Graduate Research Fellowship support. M.L. and P.L. were also partially supported by the Simons Foundation, M.L. by NIH grant R01GM108348 (to B.B.), and P.L. by NIH training grant 5T32HG004947-04. M.S. acknowledges support from the Max Planck Society. N.P., P.M., and D.R. are grateful for support from NSF HOMINID grant #1032255 and NIH grant GM100233. D.R. is an Investigator at the Howard Hughes Medical Institute.

## Author contributions

All authors contributed to the design of the study and the analysis of data. M.L. and P.L. performed the computational experiments. M.L., P.L., B.B., and D.R. wrote the manuscript with input from all authors.

## Supplementary Information for Reconstructing Austronesian population history in Island Southeast Asia

**Supplementary Figure 1.**
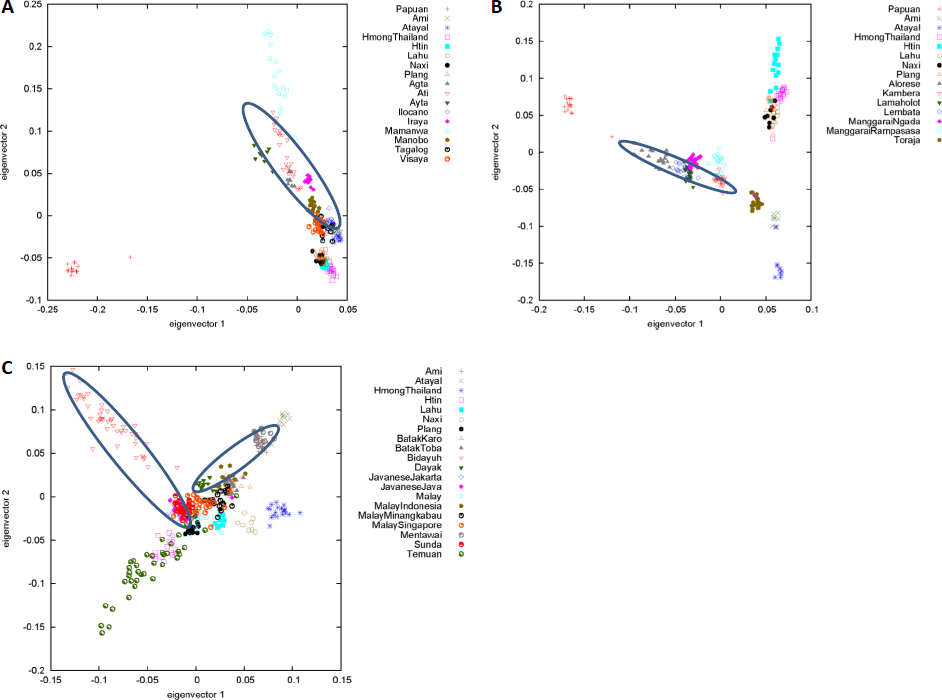
PCA plots generated with EIGENSOFT (42) for AN-speaking groups from (A) the Philippines, (B) eastern Indonesia, and (C) western ISEA, along with reference populations. The circled groupings indicate subsets of populations consistent with simple histories according to our *f*_4_-based test: (A) Agta, Ati, Ayta, Ilocano, Iraya, and Manobo (one wave of admixture), (B) Alorese, Kambera, Lamaholot, and Lembata (one wave), and (C) Bidayuh, Dayak, Mentawai, Javanese Jakarta, Javanese Java, and Sunda (two waves).

**Supplementary Figure 2.**
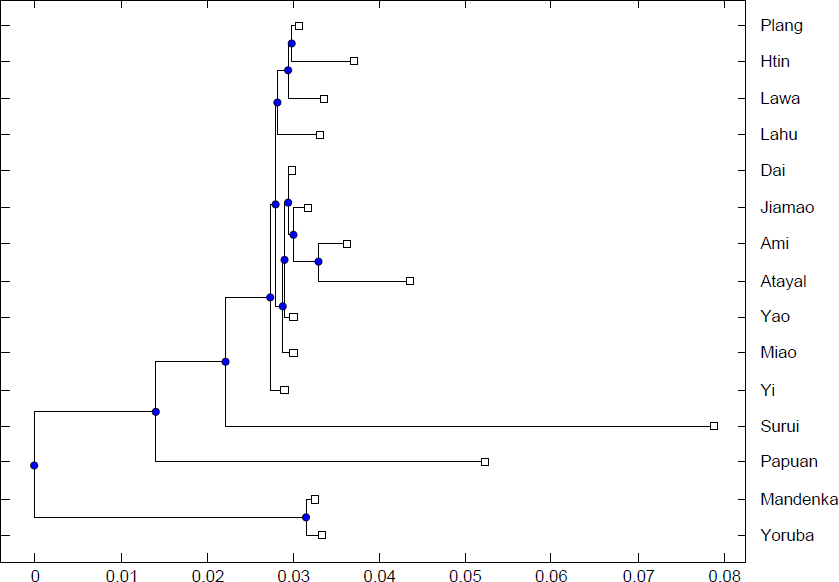
Alternative 15-population scaffold tree. See Supplementary Tables 8 and 9 for full *MixMapper* results from fitting admixed Austronesian-speaking populations using this scaffold. Distances are in *F*_2_ units.

**Supplementary Figure 3.**
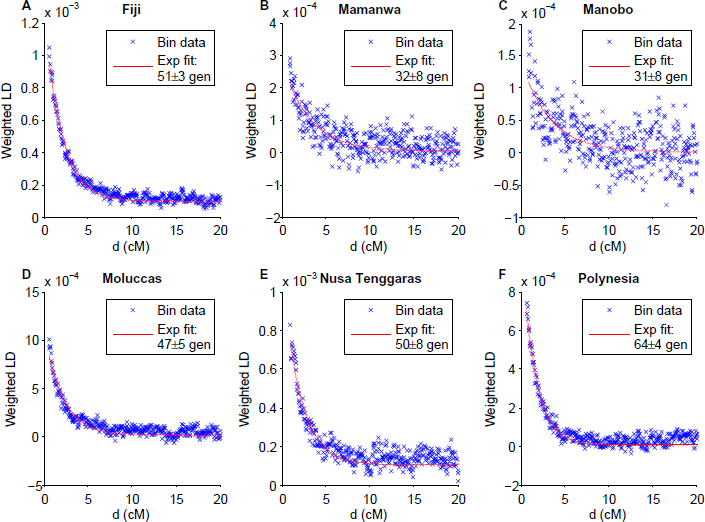
Weighted LD curves and estimated dates of admixture for (A) Fiji, (B) Mamanwa, (C) Manobo, (D) Moluccas, (E) Nusa Tenggaras, and (F) Polynesia, obtained using *ALDER* (35) with Papuan and Taiwanese reference populations. Admixture dates are inferred as time constants of the exponential decay of weighted covariance with genetic distance. LD analysis requires a higher SNP density than is available with our full data set, so these inferences are restricted to samples from ref. (27). We note that our dates are much more recent than those reported in ref. (24); we hypothesize that the initial admixtures were followed by more recent mixing between groups with different proportions of Taiwan-related ancestry, in which case the date from *ALDER* is an intermediate one over the entire process. This would be consistent with the fact that the curves appear to have some deviations from a pure exponential decay shape.

**Supplementary Figure 4.**
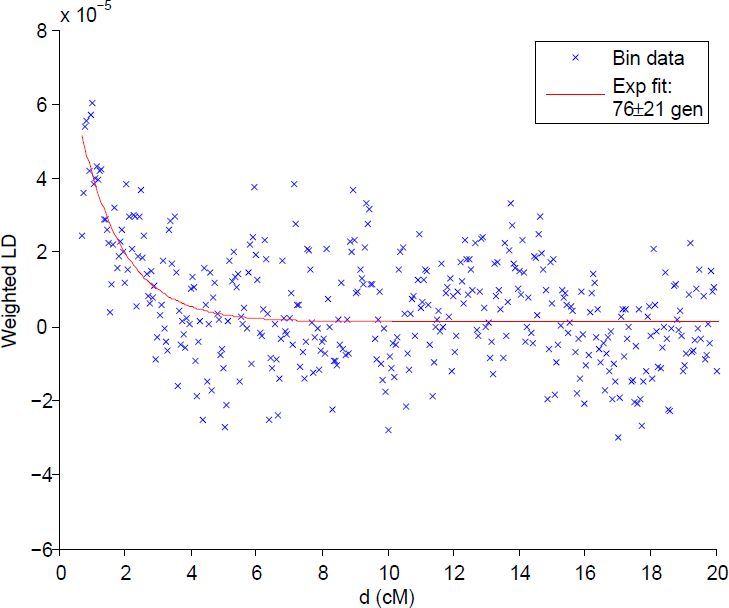
Weighted LD curve and estimated date of admixture for western ISEA, obtained using *ALDER* (35) with Papuan and CHB (HapMap Chinese from Beijing (44)) reference populations. The admixture date is inferred as the time constant of the exponential decay of weighted covariance with genetic distance. LD analysis requires a higher SNP density than is available with our full data set, so these inferences are restricted to samples from ref. (27). In order to enhance the signal-to-noise ratio, we pooled samples from four populations, two each from Borneo (Bidayuh and Dayak) and Sumatra (Besemah and Semende), into a single test set, under the assumption that all four have similar admixture histories.

**Supplementary Table 1.**
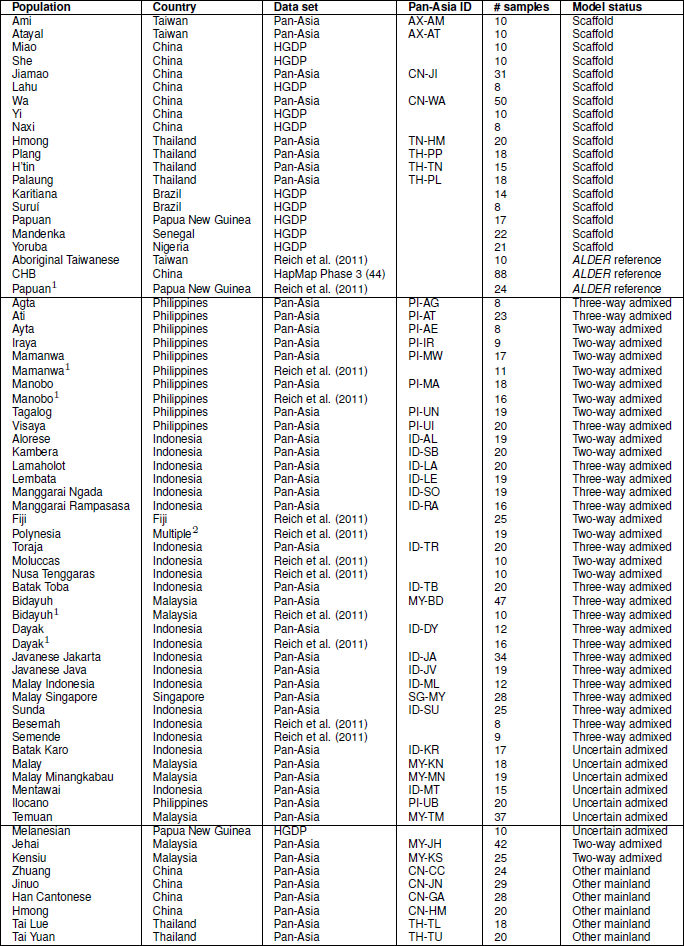
Summary of populations used in this study. The first group of populations are references used in the 18-population scaffold tree and for admixture date estimation, the second group are Austronesian-speaking populations fit as admixtures, and the third group are other populations used for comparison.

**Supplementary Table 2.** Populations with negative *f*_3_ statistics. Asian populations from Supplementary Table 1 having at least one negative *f*_3_ value. For each test population *C*, we show the two reference populations *A* and *B* in the data set giving the lowest Z-score for *f*_3_(*C*; *A, B*). We note that all populations on this list that are used in the scaffold have *Z* > −2, which indicates a non-significant result (especially given the presence of many hypotheses). While a significantly negative *f*_3_ value demonstrates that the test population must be admixed, a lack of a negative value does not prove a lack of admixture.

| Test population $C$ | Reference population $A$ | Reference population $B$ | $f_3(C; A, B)$ | Std error | Z-score |
| --- | --- | --- | --- | --- | --- |
| Alorese | Tagalog | Papuan | -0.0106 | 0.00023 | -45.36 |
| Batak Karo | Mentawai | Kalash | -0.00179 | 0.00025 | -7.21 |
| Batak Toba | Mentawai | Tuscan | -0.00238 | 0.00024 | -9.96 |
| Han Cantonese | Korean | Jiamao | -0.00079 | 0.00007 | -12.04 |
| Hmong China | Hmong Thailand | Mbuti Pygmy | -0.00132 | 0.0002 | -6.55 |
| Ilocano | Ami | Bengali | -0.00098 | 0.00024 | -4.14 |
| Javanese Jakarta | Ilocano | Jehai | -0.00113 | 0.00015 | -7.43 |
| Javanese Java | Ami | Jehai | -0.00133 | 0.00017 | -7.69 |
| Kambera | Tagalog | Papuan | -0.00719 | 0.00025 | -29.26 |
| Lamaholot | Toraja | Papuan | -0.0091 | 0.00022 | -41.2 |
| Lembata | Toraja | Papuan | -0.00961 | 0.00022 | -43.26 |
| Malay | Zhuang | GIH | -0.00322 | 0.00013 | -24.5 |
| Malay Indonesia | Ami | Bengali | -0.00201 | 0.00028 | -7.19 |
| Malay Minangkabau | Ami | Hindi Haryana | -0.00262 | 0.00026 | -9.97 |
| Malay Singapore | Hindi Haryana | Jiamao | -0.00209 | 0.00011 | -18.66 |
| Manggarai Ngada | Tagalog | Papuan | -0.00883 | 0.00025 | -35.49 |
| Manggarai Rampasasa | Ilocano | Papuan | -0.00682 | 0.00029 | -23.88 |
| Manobo | Ami | Papuan | -0.0006 | 0.00035 | -1.7 |
| Miao | Hmong Thailand | Colombian | -0.0004 | 0.00028 | -1.41 |
| Plang | Mlabri | Han-NChina | -0.00021 | 0.00027 | -0.78 |
| Sunda | Ilocano | Jehai | -0.00113 | 0.00014 | -8.15 |
| Tagalog | Ami | Hindi Rajasthan | -0.00214 | 0.00019 | -11.11 |
| Tai Yuan | Htin | CHB | -0.00082 | 0.00008 | -9.76 |
| Toraja | Ilocano | Papuan | -0.00213 | 0.00026 | -8.18 |
| Visaya | Ami | Hindi Rajasthan | -0.00298 | 0.0002 | -14.79 |
| Wa | Mlabri | Naxi | -0.00005 | 0.00028 | -0.16 |
| Yi | Mlabri | Naxi | -0.00006 | 0.00034 | -0.18 |
| Zhuang | Jiamao | Lahu | -0.0002 | 0.0001 | -1.91 |

**Supplementary Table 3.** Populations with no negative *f*_3_ statistics. Asian populations from Supplementary Table 1 having no negative *f*_3_ value for any pair of reference populations in the data set. While a significantly negative *f*_3_ value demonstrates that the test population must be admixed, a lack of a negative value does not prove a lack of admixture.

|  |
| --- |
| Agta |
| Ami |
| Atayal |
| Ati |
| Ayta |
| Bidayuh |
| Dayak |
| Hmong Thailand |
| Htin |
| Iraya |
| Jehai |
| Jiamao |
| Jinuo |
| Kensiu |
| Lahu |
| Mamanwa |
| Mentawai |
| Mlabri |
| Naxi |
| Paluang |
| She |
| Tai Lue |
| Temuan |

**Supplementary Table 4.** Inferred mixture parameters for two-way admixed populations. Sources of ancestry and mixture proportions (95% confidence intervals) from *MixMapper* for two-way admixed populations. “Papuan opp. African” refers to the common ancestral branch of all populations in the scaffold other than Papuan and Africans, while (Ami, Atayal) designates the common ancestral branch of Ami and Atayal (see Fig. 1). Branch topologies are shown that occur for at least 5% of 500 bootstrap replicates.

| **Philippine admixed population** | **Mixing branch 1 bootstrap distribution** |  | **Mixing branch 2 bootstrap distribution** |  | **Branch 1 ancestry (Austronesian)** |
| --- | --- | --- | --- | --- | --- |
| Agta | (Ami,Atayal) | 44% | Papuan | 100% | 51–62% |
|  | Ami | 56% |  |  |  |
| Ati | (Ami,Atayal) | 15% | Papuan | 100% | 50–59% |
|  | Ami | 85% |  |  |  |
| Ayta | (Ami,Atayal) | 20% | Papuan | 100% | 25–38% |
|  | Ami | 7% |  |  |  |
|  | Atayal | 73% |  |  |  |
| Iraya | (Ami,Atayal) | 28% | Papuan | 76% | 61–80% |
|  | Ami | 72% | Papuan opp. African | 20% |  |
| Mamanwa | (Ami,Atayal) | 25% | Papuan | 100% | 51–61% |
|  | Ami | 62% |  |  |  |
|  | Atayal | 13% |  |  |  |
| Manobo | (Ami,Atayal) | 11% | Papuan | 100% | 78–83% |
|  | Ami | 89% |  |  |  |
| Tagalog | (Ami,Atayal) | 99% | Papuan | 71% | 83–92% |
|  |  |  | Papuan opp. African | 28% |  |
| Visaya | (Ami,Atayal) | 88% | Papuan | 85% | 74–85% |
|  | Ami | 11% | Papuan opp. African | 15% |  |
| **E. Indonesian / Oceanian admixed population** | **Mixing branch 1 bootstrap distribution** |  | **Mixing branch 2 bootstrap distribution** |  | **Branch 1 ancestry (Austronesian)** |
| Alorese | (Ami,Atayal) | 77% | Papuan | 100% | 37–44% |
|  | Ami | 17% |  |  |  |
|  | Atayal | 6% |  |  |  |
| Fiji | (Ami,Atayal) | 19% | Papuan | 100% | 30–41% |
|  | Ami | 64% |  |  |  |
|  | Atayal | 17% |  |  |  |
| Kambera | (Ami,Atayal) | 100% | Papuan | 100% | 67–73% |
| Lamaholot | (Ami,Atayal) | 93% | Papuan | 100% | 50–56% |
|  | Ami | 6% |  |  |  |
| Lembata | (Ami,Atayal) | 94% | Papuan | 100% | 47–53% |
| Polynesia | (Ami,Atayal) | 20% | Papuan | 100% | 61–72% |
|  | Ami | 54% |  |  |  |
|  | Atayal | 26% |  |  |  |

**Supplementary Table 5.** Inferred mixture parameters for three-way admixed populations. Mixture parameters from *MixMapper* for three-way admixed populations. Mixture proportions shown are 95% confidence intervals for re-optimized values (see Methods), using the bootstrap replicates (percentages given, out of 500) assigning the third ancestry component to the H’tin branch.

| **E. Indonesian / Oceanian admixed population** | **Percent bootstrap reps with Branch 3 = H'tin** | **Branch 3 ancestry (Austro-Asiatic)** | **Branch 1 ancestry (Austronesian)** |
| --- | --- | --- | --- |
| Manggarai Ngada | 100% | 24–29% | 31–37% |
| Manggarai Rampasasa | 100% | 34–41% | 29–37% |
| Toraja | 100% | 10–17% | 68–75% |
| **W. Indonesian admixed population** | **Percent bootstrap reps with Branch 3 = H'tin** | **Branch 3 ancestry (Austro-Asiatic)** | **Branch 1 ancestry (Austronesian)** |
| Batak Toba | 92% | 22–32% | 50–57% |
| Bidayuh | 100% | 50–57% | 37–44% |
| Dayak | 100% | 35–42% | 48–56% |
| Javanese Jakarta | 100% | 57–63% | 29–35% |
| Javanese Java | 100% | 57–64% | 28–34% |
| Malay Indonesia | 100% | 26–34% | 56–64% |
| Malay Singapore | 100% | 38–45% | 37–43% |
| Sunda | 100% | 54–61% | 30–36% |

**Supplementary Table 6.** Inferred mixture parameters for two-way admixed populations with alternative SNP ascertainment. Sources of ancestry and mixture proportions (95% confidence intervals) from *MixMapper* for two-way admixed populations, using SNPs selected by merging the Pan-Asia data with HGDP samples typed on the Affymetrix Human Origins array (30). “Papuan opp. African” refers to the common ancestral branch of all populations in the scaffold other than Papuan and Africans, while (Ami, Atayal) designates the common ancestral branch of Ami and Atayal (see Fig. 1). Branch topologies are shown that occur for at least 5% of 500 bootstrap replicates. The results are very similar to those obtained with the original scaffold (see Supplementary Table 4).

| **Philippine admixed population** | **Mixing branch 1 bootstrap distribution** | **Mixing branch 2 bootstrap distribution** | **Branch 1 ancestry (Austronesian)** |
| --- | --- | --- | --- |
| Agta | (Ami,Atayal) 51%<br>Ami 38%<br>Atayal 11% | Papuan 62% | 51–66% |
| Ati | (Ami,Atayal) 93% | Papuan 100% | 53–68% |
| Ayta | (Ami,Atayal) 31%<br>Ami 17%<br>Atayal 48% | Papuan 89% | 23–45% |
| Iraya | (Ami,Atayal) 29%<br>Ami 59%<br>Atayal 12% | Papuan 35%<br>Papuan opp. African 6% | 60–86% |
| Mamanwa | (Ami,Atayal) 41%<br>Ami 42%<br>Atayal 17% | Papuan 100% | 49–66% |
| Manobo | (Ami,Atayal) 33%<br>Ami 66% | Papuan 100% | 77–87% |
| Tagalog | (Ami,Atayal) 98% | Papuan 78% | 85–93% |
| Visaya | (Ami,Atayal) 87%<br>Ami 5%<br>Atayal 8% | Papuan 99% | 82–91% |
| **E. Indonesian / Oceanian admixed population** | **Mixing branch 1 bootstrap distribution** | **Mixing branch 2 bootstrap distribution** | **Branch 1 ancestry (Austronesian)** |
| Alorese | (Ami,Atayal) 72%<br>Atayal 20% | Papuan 100% | 38–47% |
| Kambera | (Ami,Atayal) 95% | Papuan 100% | 65–75% |
| Lamaholot | (Ami,Atayal) 93% | Papuan 100% | 51–62% |
| Lembata | (Ami,Atayal) 55%<br>Ami 15%<br>Atayal 30% | Papuan 100% | 48–57% |

**Supplementary Table 7.** Inferred mixture parameters for three-way admixed populations with alternative SNP ascertainment. Mixture parameters from *MixMapper* for three-way admixed populations, using SNPs selected by merging the Pan-Asia data with HGDP samples typed on the Affymetrix Human Origins array (30). Mixture proportions shown are 95% confidence intervals for re-optimized values (see Methods), using the bootstrap replicates (percentages given, out of 500) assigning the third ancestry component to the H’tin branch. The results are very similar to those obtained with the original scaffold (see Supplementary Table 5), with slightly lower but still substantial bootstrap support for the H’tin-related ancestry component.

| **E. Indonesian / Oceanian admixed population** | **Percent bootstrap reps with Branch 3 = H'tin** | **Branch 3 ancestry (Austro-Asiatic)** | **Branch 1 ancestry (Austronesian)** |
| --- | --- | --- | --- |
| Manggarai Ngada | 66% | 20–31% | 29–42% |
| Manggarai Rampasasa | 27% | 29–38% | 33–44% |
| Toraja | 85% | 6–14% | 70–79% |
| **W. Indonesian admixed population** | **Percent bootstrap reps with Branch 3 = H'tin** | **Branch 3 ancestry (Austro-Asiatic)** | **Branch 1 ancestry (Austronesian)** |
| Batak Toba | 28% | 19–35% | 49–60% |
| Bidayuh | 99% | 42–58% | 36–50% |
| Dayak | 98% | 27–44% | 46–59% |
| Javanese Jakarta | 100% | 49–64% | 28–40% |
| Javanese Java | 100% | 52–70% | 24–38% |
| Malay Indonesia | 76% | 18–33% | 58–73% |
| Malay Singapore | 74% | 29–49% | 35–51% |
| Sunda | 100% | 50–65% | 27–41% |

**Supplementary Table 8.** Inferred mixture parameters for two-way admixed populations on a 15-population alternative scaffold. Sources of ancestry and mixture proportions (95% confidence intervals) from *MixMapper* for two-way admixed populations using a 15-population alternative scaffold tree. The results are very similar to those obtained with the original scaffold (see Supplementary Table 4). “Papuan opp. African” refers to the common ancestral branch of all populations in the scaffold other than Papuan and Africans, while (Ami, Atayal) designates the common ancestral branch of Ami and Atayal (see Fig. 1). Branch topologies are shown that occur for at least 5% of 500 bootstrap replicates.

| **Philippine admixed population** | **Mixing branch 1 bootstrap distribution** |  | **Mixing branch 2 bootstrap distribution** |  | **Branch 1 ancestry (Austronesian)** |
| --- | --- | --- | --- | --- | --- |
| Agta | (Ami,Atayal) | 44% | Papuan | 100% | 50–60% |
|  | Ami | 56% |  |  |  |
| Ati | (Ami,Atayal) | 12% | Papuan | 100% | 49–58% |
|  | Ami | 88% |  |  |  |
| Ayta | (Ami,Atayal) | 21% | Papuan | 100% | 24–37% |
|  | Ami | 7% |  |  |  |
|  | Atayal | 72% |  |  |  |
| Iraya | (Ami,Atayal) | 16% | Papuan | 39% | 56–78% |
|  | Ami | 84% | Papuan opp. African | 60% |  |
| Mamanwa | (Ami,Atayal) | 40% | Papuan | 100% | 51–61% |
|  | Ami | 53% |  |  |  |
|  | Atayal | 7% |  |  |  |
| Manobo | (Ami,Atayal) | 9% | Papuan | 100% | 78–83% |
|  | Ami | 91% |  |  |  |
| Tagalog | (Ami,Atayal) | 100% | Papuan | 64% | 83–92% |
|  |  |  | Papuan opp. African | 34% |  |
| Visaya | (Ami,Atayal) | 82% | Papuan | 78% | 72–85% |
|  | Ami | 18% | Papuan opp. African | 22% |  |
| **E. Indonesian / Oceanian admixed population** | **Mixing branch 1 bootstrap distribution** |  | **Mixing branch 2 bootstrap distribution** |  | **Branch 1 ancestry (Austronesian)** |
| Alorese | (Ami,Atayal) | 84% | Papuan | 100% | 37–43% |
|  | Ami | 14% |  |  |  |
| Fiji | (Ami,Atayal) | 16% | Papuan | 100% | 30–40% |
|  | Ami | 66% |  |  |  |
|  | Atayal | 18% |  |  |  |
| Kambera | (Ami,Atayal) | 100% | Papuan | 100% | 68–72% |
| Lamaholot | (Ami,Atayal) | 96% | Papuan | 100% | 49–56% |
| Lembata | (Ami,Atayal) | 98% | Papuan | 100% | 47–53% |
| Polynesia | (Ami,Atayal) | 23% | Papuan | 100% | 61–72% |
|  | Ami | 52% |  |  |  |
|  | Atayal | 25% |  |  |  |

**Supplementary Table 9.**
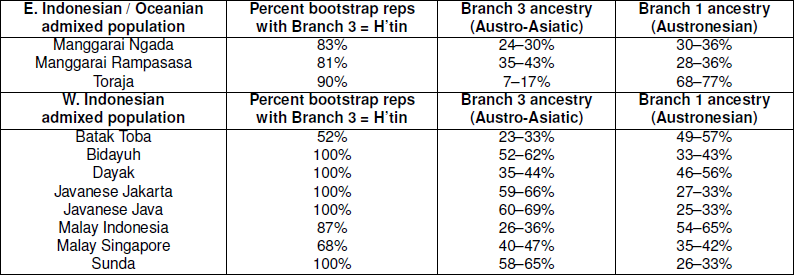
Inferred mixture parameters for three-way admixed populations on a 15-population alternative scaffold. Mixture parameters from *MixMapper* for three-way admixed populations using a 15-population alternative scaffold tree. The results are very similar to those obtained with the original scaffold (see Supplementary Table 5), with slightly lower but still substantial bootstrap support for the H’tin-related ancestry component. Mixture proportions shown are 95% confidence intervals for re-optimized values (see Methods), using the bootstrap replicates (percentages given, out of 500) assigning the third ancestry component to the H’tin branch.

**Supplementary Table 10.** Consistency of mixture parameters for two-way admixed populations on 17-population alternative scaffolds. Sources of ancestry and mixture proportions (95% confidence intervals) from *MixMapper* for two-way admixed populations, removing one population at a time (other than Papuan) from the 18-population scaffold tree (Fig. 1). Values are means ± standard errors over the 17 different perturbed scaffolds. Austronesian ancestry refers to splits from the Ami and Atayal branches and their common ancestor, while Papuan support only includes splits from the Papuan branch. The results are very similar to those obtained with the original scaffold (Supplementary Table 4). Note that the branch support values are over 100 replicates, while the mixture proportions are point-estimates using all data rather than bootstraps.

| **Philippine admixed population** | **Taiwan bootstrap support** | **Papuan bootstrap support** | **Taiwan ancestry fraction** |
| --- | --- | --- | --- |
| Agta | 100 ± 0% | 99 ± 0% | 56 ± 1% |
| Ati | 100 ± 0% | 100 ± 0% | 55 ± 0% |
| Ayta | 99 ± 1% | 100 ± 0% | 32 ± 1% |
| Iraya | 100 ± 0% | 79 ± 8% | 73 ± 2% |
| Mamanwa | 100 ± 0% | 100 ± 0% | 56 ± 0% |
| Manobo | 100 ± 0% | 100 ± 0% | 81 ± 0% |
| Tagalog | 100 ± 0% | 69 ± 11% | 89 ± 0% |
| Visaya | 100 ± 0% | 83 ± 6% | 83 ± 0% |
| **E. Indonesian / Oceanian admixed population** | **Taiwan bootstrap support** | **Papuan bootstrap support** | **Taiwan ancestry fraction** |
| Alorese | 100 ± 0% | 100 ± 0% | 40 ± 0% |
| Fiji | 100 ± 1% | 100 ± 0% | 36 ± 0% |
| Kambera | 100 ± 0% | 100 ± 0% | 70 ± 1% |
| Lamaholot | 100 ± 0% | 100 ± 0% | 53 ± 1% |
| Lembata | 100 ± 0% | 100 ± 0% | 50 ± 1% |
| Polynesia | 100 ± 0% | 100 ± 0% | 66 ± 0% |

**Supplementary Table 11.** Consistency of mixture parameters for three-way admixed populations on 17-population alternative scaffolds. Mixture parameters from *MixMapper* for three-way admixed populations, removing one population at a time (other than Papuan) from the 18-population scaffold tree (Fig. 1). Values are means ± standard errors over the 17 different perturbed scaffolds. The results are very similar to those obtained with the original scaffold (see Supplementary Table 5). Mixture proportions shown are re-optimized values (see Methods), using the 17-population trees in which the third ancestry component is Austro-Asiatic (H’tin, Plang, or Wa), which were 16 of 17 for Batak Toba and Manggarai Rampasasa and all 17 trees for the other populations. Note that the branch support values are over 100 replicates, while the mixture proportions are point-estimates using all data rather than bootstraps.

| **E. Indonesian / Oceanian admixed population** | **Percent bootstrap reps with Branch 3 = H'tin, Plang, Wa** | **Branch 3 ancestry (Austro-Asiatic)** | **Branch 1 ancestry (Austronesian)** |
| --- | --- | --- | --- |
| Manggarai Ngada | 100 ± 1% | 26 ± 2% | 34 ± 2% |
| Manggarai Rampasasa | 96 ± 15% | 37 ± 2% | 34 ± 2% |
| Toraja | 99 ± 4% | 12 ± 1% | 72 ± 1% |
| **W. Indonesian admixed population** | **Percent bootstrap reps with Branch 3 = H'tin, Plang, Wa** | **Branch 3 ancestry (Austro-Asiatic)** | **Branch 1 ancestry (Austronesian)** |
| Batak Toba | 93 ± 12% | 27 ± 2% | 53 ± 2% |
| Bidayuh | 100 ± 0% | 54 ± 2% | 40 ± 1% |
| Dayak | 100 ± 0% | 39 ± 2% | 52 ± 1% |
| Javanese Jakarta | 100 ± 0% | 59 ± 2% | 32 ± 2% |
| Javanese Java | 100 ± 0% | 60 ± 2% | 31 ± 2% |
| Malay Indonesia | 97 ± 9% | 31 ± 1% | 60 ± 1% |
| Malay Singapore | 98 ± 7% | 41 ± 3% | 40 ± 2% |
| Sunda | 100 ± 0% | 57 ± 2% | 33 ± 1% |

**Supplementary Table 12.** Admixture model selection for three-way admixed populations. Quality of fit for alternative models for three-way admixed populations. Shown are the median norms of the vectors of residual errors for all pairwise distances *f*_2_(*C, X*) (see Methods for details), along with 95% confidence intervals for the differences (all multiplied by 10^6^). Smaller norms indicate more accurate model fits.

| **E. Indonesian / Oceanian admixed population** | **Residual norm from 2-way fit** | **Residual norm from 3-way fit** | **Difference (95% CI)** |
| --- | --- | --- | --- |
| Manggarai Ngada | 27.0 | 22.7 | (-1.4, 9.8) |
| Manggarai Rampasasa | 31.2 | 25.1 | (-1.4, 14.5) |
| Toraja | 11.3 | 7.9 | (-0.8, 7.0) |
| **W. Indonesian admixed population** | **Residual norm from 2-way fit** | **Residual norm from 3-way fit** | **Difference (95% CI)** |
| Batak Toba | 22.2 | 16.5 | (-5.5, 15.2) |
| Bidayuh | 23.1 | 15.5 | (-1.6, 16.5) |
| Dayak | 32.8 | 11.1 | (11.4, 28.1) |
| Javanese Jakarta | 34.3 | 15.3 | (12.4, 23.8) |
| Javanese Java | 32.8 | 15.0 | (10.8, 24.0) |
| Malay Indonesia | 18.8 | 10.1 | (0.8, 14.9) |
| Malay Singapore | 38.8 | 27.0 | (0.6, 21.1) |
| Sunda | 39.1 | 16.8 | (15.8, 27.8) |

**Supplementary Table 13.** Two-way mixture fits for East and Mainland Southeast Asian populations. Inferred sources of ancestry (with bootstrap support) and mixture proportions (95% confidence intervals) from *MixMapper* for East and Mainland Southeast Asian populations. Names with parentheses refer to the common ancestral branches of the specified nodes (see Fig. 1). Branch topologies are shown that occur for at least 5% of 500 bootstrap replicates. We see essentially no evidence of the four ancestry components found in Austronesian-speaking groups, aside from H’tin-related (Austro-Asiatic) ancestry in several populations. We note that some of the populations here may not truly be admixed, but we show all of the fits for completeness.

| Admixed population | Mixing branch 1 + branch 2 | % reps | Branch 1 ancestry |
| --- | --- | --- | --- |
| Chinese Singapore | (Ami,Atayal,Jiamao) + Karitiana | 56% | 98–99% |
|  | (Ami,Atayal,Jiamao) + Naxi | 21% | 85–93% |
|  | (Ami,Atayal,Jiamao) + Surui | 15% | 98–100% |
| Han Hakka | (Ami,Atayal,Jiamao) + Naxi | 75% | 83–91% |
|  | (Ami,Atayal,Jiamao) + She | 9% | 58–89% |
| Han Minnan | (Ami,Atayal,Jiamao) + Naxi | 63% | 84–91% |
|  | (Ami,Atayal,Jiamao) + Surui | 13% | 99–99% |
|  | (Ami,Atayal,Jiamao) + Karitiana | 13% | 99–99% |
|  | (Ami,Atayal,Jiamao) + She | 8% | 60–88% |
| Hmong China | Hmong Thailand + Jiamao | 40% | 71–89% |
|  | Hmong Thailand + (Ami,Atayal,Jiamao) | 34% | 57–74% |
|  | Hmong Thailand + She | 20% | 56–80% |
| Jinuo | (H'tin,Plang,Wa) + Yi | 16% | 77–91% |
|  | (Naxi,Yi) + Wa | 12% | 52–80% |
|  | (Karitiana,Mandenka,Naxi,Papuan,Surui,Yi,Yoruba,root) + Wa | 11% | 65–88% |
|  | (H'tin,Plang,Wa) + (Naxi,Yi) | 8% | 41–83% |
|  | (H'tin,Plang,Wa) + Hmong Thailand | 7% | 82–97% |
|  | (H'tin,Plang,Wa) + Papuan | 7% | 97–99% |
|  | (H'tin,Plang,Wa) + Naxi | 6% | 74–93% |
| Karen | (H'tin,Plang,Wa) + Papuan | 93% | 92–98% |
|  | (H'tin,Plang) + Papuan | 7% | 90–96% |
| Lawa | (H'tin,Plang) + Papuan | 82% | 93–98% |
|  | (H'tin,Plang,Wa) + Papuan | 5% | 95–98% |
|  | H'tin + Papuan | 5% | 93–98% |
| Mlabri | H'tin + Papuan | 70% | 86–97% |
|  | H'tin + (Mandenka,Yoruba,root) | 18% | 85–95% |
|  | H'tin + (Mandenka,Yoruba) | 9% | 92–98% |
| Mon | (H'tin,Plang,Wa) + (Mandenka,Yoruba,root) | 90% | 80–86% |
| Tai Khuen | Jiamao + H'tin | 99% | 65–75% |
| Tai Lue | Jiamao + H'tin | 97% | 68–81% |
| Tai Yong | Jiamao + H'tin | 95% | 66–76% |
| Tai Yuan | Jiamao + H'tin | 86% | 48–60% |
|  | (Ami,Atayal,Jiamao) + H'tin | 10% | 56–66% |
| Yao | (Ami,Atayal,Jiamao) + Hmong Thailand | 79% | 60–86% |
|  | Hmong Thailand + H'tin | 6% | 87–94% |
|  | (Ami,Atayal,Jiamao,She) + Hmong Thailand | 6% | 81–89% |
| Zhuang | Jiamao + H'tin | 99% | 87–92% |

**Supplementary Table 14.** Formal test for numbers of sources of admixture. We applied a formal test based on *f*_4_ statistics, as described in refs. (33) and (34), to estimate how many sources of admixture are necessary to explain the observed relationships among a collection of admixed populations. Briefly, we estimate the rank of a matrix of values *f*_4_(*A, B*; *C, D*), where *A* and *B* are populations in a test set and *C* and *D* are populations in a reference set. To remove trivially linearly dependent rows and columns, we fix *A* and *C* to be the first populations in each list (without loss of generality) and let *B* and *D* vary. In order to maximize sensitivity for separate sources of Asian ancestry, we used a reference set consisting of Yoruba as the fixed outgroup *C* and 31 East and Southeast Asian populations as the other references *D*. We used a *p*-value threshold of 0.05; a score below this threshold implies that at least that many sources are necessary to explain the relationships among the test set. In bold are the maximal significant values, indicating the estimated number of sources for each set.

| Test subset | *p*-value for 2 sources | *p*-value for 3 sources | *p*-value for 4 sources |
| --- | --- | --- | --- |
| Agta, Ati, Ayta, Ilocano, Iraya, Manobo | **0.000** | 0.110 | 0.156 |
| Alorese, Kambera, Lamaholot, Lembata | **0.000** | 0.486 | 0.428 |
| Alorese, Kambera, Lamaholot, Lembata, Manggarai Ngada, Manggarai Rampasasa | 0.000 | **0.000** | 0.366 |
| Bidayuh, Dayak, Javanese Jakarta, Javanese Java, Mentawai, Sunda | 0.000 | **0.000** | 0.068 |
| Bidayuh, Dayak, Javanese Jakarta | 0.000 | **0.018** | NA |

**Supplementary Table 15.** Mixture fits for Austronesian-speaking populations with no Taiwanese in the scaffold tree. Inferred sources of ancestry (with bootstrap support) and mixture proportions (95% confidence intervals) from *MixMapper* for selected Austronesian-speaking populations, using a 16-population scaffold tree formed by removing Ami and Atayal from the original scaffold (i.e., Miao, She, Jiamao, Lahu, Wa, Yi, Naxi, Hmong, Plang, H’tin, Palaung, Karitiana, Suruí, Papuan, Mandenka, and Yoruba). Names with parentheses refer to the common ancestral branches of the specified nodes (see Fig. 1). Branch topologies are shown that occur for at least 5% of 500 bootstrap replicates. We report admixture fits for Ami and Atayal as test populations, as well as all other Austronesian-speaking populations with no negative *f*_3_ statistics (Supplementary Table 3) and selected others to fill in geographic coverage gaps. For both Ami and Atayal, more than half of the bootstrap replicates yield fits with 90% or more Jiamao ancestry and a very small proportion of a seemingly implausible second ancestry component (e.g., Native American). In our experience, such results indicate that the test populations should in fact be modeled as unadmixed relative to the scaffold, in this case adjacent to Jiamao (31). For other populations, meanwhile, the fits appear to be reasonable and are very similar (both in topology and mixture proportions) to those obtained with the original scaffold (with the difference that Jiamao is now the closest population to the previous location of the Taiwanese). Fits with Manobo reported as one mixing branch are three-way admixtures (proportions are not re-optimized).

| Admixed population | Mixing branch 1 + branch 2 | % reps | Branch 1 ancestry |
| --- | --- | --- | --- |
| Agta | Jiamao + Papuan | 96% | 51–62% |
| Alorese | Papuan + H'tin | 81% | 55–62% |
|  | Papuan + (H'tin,Plang) | 9% | 55–62% |
| Ami | Jiamao + H'tin | 43% | 85–95% |
|  | Jiamao + Karitiana | 36% | 98–99% |
|  | Jiamao + Surui | 15% | 98–99% |
| Atayal | Jiamao + (Karitiana,Surui) | 30% | 88–97% |
|  | Jiamao + Papuan | 14% | 93–99% |
|  | Jiamao + (Mandenka,Papuan,Yoruba,root) | 12% | 88–97% |
|  | Jiamao + (Karitiana,Mandenka,Papuan,Surui,Yoruba,root) | 10% | 71–88% |
|  | Jiamao + (Mandenka,Yoruba,root) | 8% | 94–98% |
|  | Jiamao + H'tin | 7% | 78–95% |
|  | Jiamao + Surui | 7% | 97–99% |
| Ati | Jiamao + Papuan | 95% | 50–59% |
| Ayta | Papuan + H'tin | 92% | 61–75% |
|  | Papuan + Jiamao | 5% | 63–80% |
| Bidayuh | Manobo + H'tin | 100% | 30–42% |
| Dayak | Manobo + H'tin | 100% | 46–59% |
| Iraya | Jiamao + Papuan | 77% | 68–80% |
|  | Jiamao + (Mandenka,Papuan,Yoruba,root) | 21% | 57–71% |
| JavaneseJakarta | Manobo + H'tin | 100% | 34–46% |
| Kambera | (H'tin,Plang,Wa) + Papuan | 42% | 71–76% |
|  | Jiamao + Papuan | 34% | 67–74% |
|  | (HmongThailand,Jiamao,Miao,She) + Papuan | 21% | 69–75% |
| Mamanwa | Jiamao + Papuan | 76% | 50–60% |
|  | H'tin + Papuan | 12% | 52–59% |
|  | (H'tin,Plang,Wa) + Papuan | 9% | 53–64% |
| Manobo | Jiamao + Papuan | 100% | 77–84% |
| Sunda | Manobo + H'tin | 100% | 40–51% |

**Supplementary Table 16.** Robustness of Austro-Asiatic ancestry with modified scaffolds. Robustness of the Austro-Asiatic ancestry component from *MixMapper* for three-way admixed populations with either H’tin or H’tin and Plang removed from the 18-population scaffold tree. Shown are the percentages of bootstrap replicates (out of 500) assigning the third ancestry component in a three-way admixture model to an Austro-Asiatic branch in the scaffold (Plang or Wa in the first column and Wa in the second column). The fits on the reduced scaffolds are not as robust for the eastern Indonesian populations, while the lower confidences for Batak Toba and the Malay populations may be due to a small proportion of Indian ancestry (20, 25) that is picked up more often with fewer Austro-Asiatic references present.

| **E. Indonesian / Oceanian admixed population** | **Percent bootstrap support with H'tin removed** | **Percent bootstrap support with H'tin and Plang removed** |
| --- | --- | --- |
| Manggarai Ngada | 95% | 16% |
| Manggarai Rampasasa | 44% | 0% |
| Toraja | 84% | 36% |
| **W. Indonesian admixed population** | **Percent bootstrap support with H'tin removed** | **Percent bootstrap support with H'tin and Plang removed** |
| Batak Toba | 45% | 24% |
| Bidayuh | 100% | 98% |
| Dayak | 100% | 93% |
| Javanese Jakarta | 100% | 100% |
| Javanese Java | 100% | 100% |
| Malay Indonesia | 69% | 29% |
| Malay Singapore | 63% | 31% |
| Sunda | 100% | 100% |

